# A stochastic model of cortical microtubule anchoring and mechanics provides regulatory control of microtubule shape

**DOI:** 10.1101/2022.09.08.507191

**Authors:** Tim Y.Y. Tian, Colin B. Macdonald, Eric N. Cytrynbaum

## Abstract

The organization of cortical microtubule arrays play an important role in the development of plant cells. Until recently, the direct mechanical influence of cell geometry on the constrained microtubule (MT) trajectories have been largely ignored in computational models. Modelling MTs as thin elastic rods constrained on a surface, a previous study examined the deflection of MTs using a fixed number of segments and uniform segment lengths between MT anchors. It is known that the resulting MT curves converge to geodesics as the anchor spacing approaches zero. In the case of long MTs on a cylinder, buckling was found for transverse trajectories. There is a clear interplay between two factors in the problem of deflection: curvature of the membrane and the lengths of MT segments. We examine the latter in detail, in the backdrop of a circular cylinder. In reality, the number of segments are not predetermined and their lengths are not uniform. We present a minimal, realistic model treating the anchor spacing as a stochastic process and examine the net effect on deflection. We find that, by tuning the ratio of growth speed to anchoring rate, it is possible to mitigate MT deflection and even prevent buckling for lengths significantly larger than the previously derived critical buckling length. We suggest that this mediation of deflection by anchoring might provide cells with a means of preventing arrays from deflecting away from the transverse orientation.

## 1 Introduction

Plant morphology on the macroscopic scale is influenced by microscopic processes within individual cells. One aspect is the formation of parallel cortical microtubule (MT) arrays. The orientation of these arrays have been shown to play a role in the deposition of cellulose in a parallel manner, with the resulting cell wall constraining the direction of cell elongation [1]. The process of MT organization is of particular interest because higher-plant cells do not have MT organization centres; instead, these MT arrays rely on self-organization. There have been many studies into the various factors which come into play: MT-MT interactions, localization of MT-related proteins, and nucleation distribution. MTs are dynamic, and their interactions with one another include zippering, catastrophe, crossover, and severing [2]. The regulation of MT-related proteins influence the attachment of MTs on the membrane and edge-induced catastrophe [3, 4].

In tandem with experiments, this process has been studied in mathematical and computational models and analyzed in simulations to understand the importance of each factor. Simulations have been run on geometries with varying levels of complexity: connected planes representing a rectangular prism [4], cylindrical prisms [5, 6], triangular meshes approximating cells [7], and within volumes of smooth solids [8]. In many of these works, the assumption is made that MTs travel along geodesics; this is justified if MT segments lengths between anchors are very short relative to the radii of curvature of the cell surface. The possibility of deflection has been studied in vitro within microchambers [9] and compared to a model where MTs are considered as thin elastic rods constrained to the membrane surface. It was found that the deflection of grown MTs, although not completely constrained to the surface of the microchamber, indeed matches that of longitudinal coiling predicted by theory – a finding inconsistent with the transverse arrays found in cells. The authors in [9] speculated that one explanation was due to unanchored lengths being very small *in vivo* by comparison to their experiments.

Recently, the growth of constrained MTs has been examined closer in theory while incorporating the effect of uniform MT segment lengths between anchoring points [10]. It was found, in the case of a circular cylinder, deflection may be significant when given uniform segments approximated to be those observed in a previous experiment; one cannot conclusively rule out the influence of elastic-rod mechanics. The different models are shown in Figure 1.

**Figure 1:**
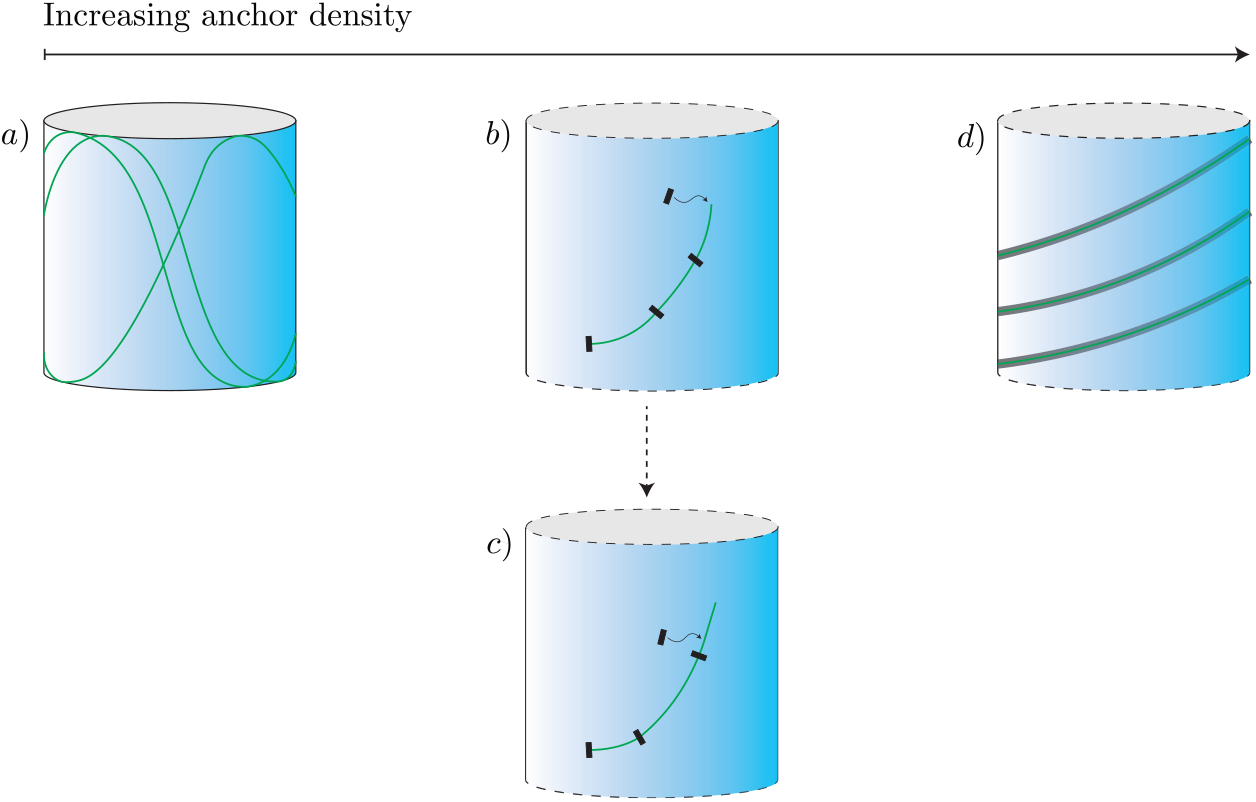
Iterations in the modelling of MT paths constrained on the cell membrane: a) MTs spanning length scales comparable to that of the cell circumference prior to anchoring in the case on inaccessible end caps [9], b) MTs with shorter segment lengths and uniform anchoring [10], c) MTs with a stochastic anchoring process examined in the present model, d) MTs with infinite anchoring density following geodesics [5, 7]. Cases b)-d) are assumed on an infinite cylinder.

The question remained on how the segment lengths are distributed in the true anchoring process and the implications this may have. The anchoring process plays a key role: it determines the extent to which elastic curvature minimization is local, as in the extreme case of geodesics, or non-local. We make the same biophysical assumptions with regards to the MT-membrane interaction and MT mechanics as in the previous study [10], described in Section 2.1, and replace the oversimplified anchoring process (in the remainder of Section 2) with an improved yet still simple model for MT-membrane anchoring. This anchoring process generates a distribution of segment lengths. The model is set in the static cylindrical geometry, and is dependent on the speed of polymerization in relation to the anchoring rate. By examining the dependence on our model parameters in Section 3, we determine the extent to which elastic-induced deflection in the context of cell-surface curvature may be reduced in comparison with the results where anchoring was treated in a simpler way.

## 2 Model

The process of MT anchoring to the membrane plays a key role: it determines the extent to which elastic curvature minimization is local, as in the extreme case of geodesics, or non-local. We make the same biophysical assumptions with regards to the MT-membrane interaction and MT mechanics as in the previous study [10], described in Section 2.1, and replace the oversimplified anchoring process (in the remainder of Section 2) with an improved yet still simple model for MT-membrane anchoring. This anchoring process generates a distribution of segment lengths. The model is set in the static cylindrical geometry, and is dependent on the speed of polymerization in relation to the anchoring rate. By examining the dependence on our model parameters in Section 3, we determine the extent to which elastic-induced deflection in the context of cell-surface curvature may be reduced in comparison with the results where anchor was treated in a simpler way.

### 2.1 Curvature Minimization

We model each free tip as an elastic rod constrained to a cylindrical membrane, with the preceding anchor determining one end of the elastic and the MT plus end determines the other. The nucleation process is assumed to determine an initial direction, and there are no constraints made on the plus-end end point other than being constrained on the surface. We assume that the path taken minimizes the total curvature energy found by integrating the curvature along the path. Due to curvature being the only energy term, the minimizing solution is independent of Young’s modulus. We parametrize trajectories by arc-length *s* ∈ [0, *L*], where length has been non-dimensionalized by the radius of the cylinder *R*. On the cylinder, it is possible to explicitly find the parametrization of a trajectory in terms of the angle *φ*(*s*) between the circumferential and longitudinal tangent directions. In cylindrical coordinates characterized with radial angle and cylinder height (*θ, z*), *φ* measures the angle from the *θ* axis. See [10] for a derivation. As a result, the energy functional is given by,

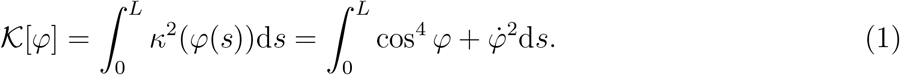

Using variational methods, we derived the Euler-Lagrange (EL) equation

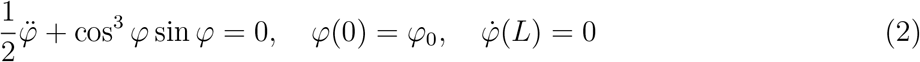

where *φ*_0_ is the initial direction of growth prescribed by the preceding anchor. The last condition arises from natural boundary conditions due to the unconstrained end point. Paths which solve Equation (2) minimize the functional and are assumed to be the shape taken on by the MT.

Without loss of generality, we restrict to trajectories in which *φ*_0_ ∈ [0, *π/*2]. Trajectories with *φ*_0_ = 0 exhibit buckling at non-dimensional length

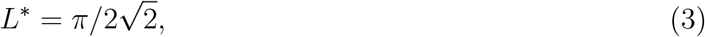

where the path *φ* = 0, stable for *L < L*^∗^, becomes unstable and is replaced by a non-zero path as a stable solution [10]. Due to the symmetry and interest in energetically favorable paths, we may further restrict to *φ*(*s*) ∈ [0, *π/*2]. Although there are stable branches of *φ*(*s*) which dip below 0 for *φ*_0_ *>* 0, these are not energetically favorable and are difficult to reach from a typical growing configuration.

### 2.2 Anchoring Times

In the following discussion, in contrast with Section 2.1, we use dimensional quantities. We assume that anchoring consists of linking MTs with the membrane by surrounding proteins. The specific proteins involved have yet to be conclusively identified, and we do not speculate on the identity. CLASP is suspected to play a role in this process [11, 3].

It is also possible that anchoring occurs at specific sites facilitated by various cellular structures. However, these are unlikely to be the only anchor sites [12]. As a simplification, we assume anchors are present uniformly in the cytosol, such that anchoring occurs at a rate proportional to the instantaneous free-tip length with rate constant *k*_on_. Furthermore, we assume anchoring may occur anywhere on the MT and be linked to any location along the membrane. The detachment of anchors is omitted from this model with the justification below.

The trajectory of the MT is determined by the free tip so the binding kinetics is only of interest at this tip. Denoting the anchoring protein and free tip as *A* and *MT*_*tip*_, respectively, the reaction kinetics are described as,

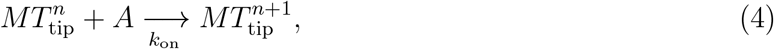

for 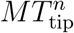 denoting the tip at the *n*th anchoring. Given tip length 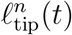, the instantaneous attachment rate is 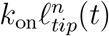. The MT growth rate is further assumed to be a constant, *v*_*g*_. Then, starting at time *t*_0_ = 0 and initial tip-length *𝓁*_0_, anchoring is an inhomogeneous Poisson process with rate *k*_on_(*𝓁*_0_ + *v*_*g*_*t*). The resulting probability density of anchoring times conditioned on *l*_0_ is

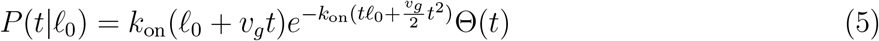

where Θ(*t*) is the Heaviside function. With a change of variables, the probability density of the length grown prior to anchoring, *𝓁*, conditioned on initial length *𝓁*_0_ is given by letting 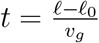:

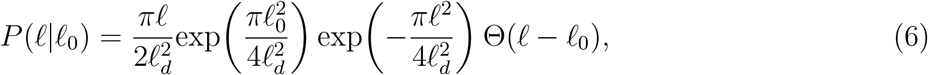

where we have introduced a length scale parameter

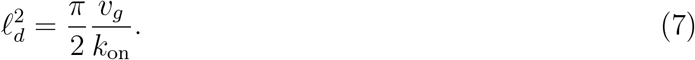

The inclusion of the *π/*2 in the definition of *𝓁*_*d*_ is for convenience, ensuring that the mean length comes out to be *𝓁*_*d*_ for the case of *𝓁*_0_ = 0.

This result is equivalent to a previous model [11] where *k*_off_*/k*_on_ → 0. In the aforementioned work [11], the detachment constant is found to be extremely small in experiments. Furthermore, the detachment only affects the MT tip trajectory if the furthest anchor falls off. Combined, these factors suggest that detachment does not affect the path significantly.

When used to determine deflection in the EL equation, we must non-dimensionalize length by *R*. Therefore, we introduce the non-dimensional length scale,

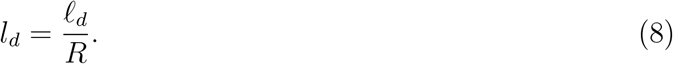

In the following sections, we use only non-dimsenionalize length variables and parameters. To distinguish these, we let the print ell (*l*) variables be non-dimensional whereas cursive ell (*𝓁*) variables are dimensional as above. Growth speed is rescaled to 1 such that time is rescaled as,

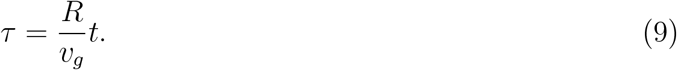

### 2.3 Event Times

For this simulation, we sample from this distribution using inverse transform sampling. Let *u* be uniformly distributed on [0, 1] and Δ*τ* be the time elapsed between anchoring events:

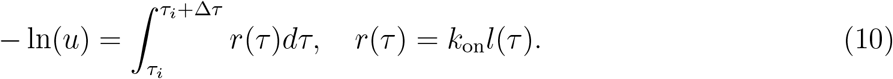

We solve the quadratic for *τ* and take the smallest root *>* 0:

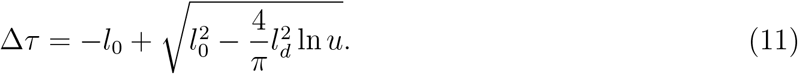

In particular, change in length before anchoring is,

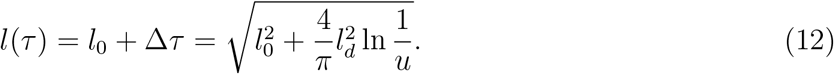

### 2.4 Anchoring Position

Consider the process starting from the nucleation anchor of a MT. After the elapsed time for the next anchoring event Δ*τ* is found, the tip length is *l*_tip_(Δ*τ*) = *l*_0_ + Δ*τ* (recall the growth speed has been normalized) and the position of anchoring *s*′ must be determined. We make the assumption that there is no preferred bonding site along MTs and let the position be uniformly distributed along the tip length, *s*′ ∼ Unif [0, *l*_tip_(Δ*τ*)]. Denoting *φ*_0_ as the initial angle, measured at the last anchor, we solve the EL equation and find 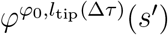: the solution with initial angle *φ*_0_ and final length *l*_tip_(Δ*τ*) evaluated at *s*′ as measured from the last anchor. After each anchoring of the tip, we shift the arc-length coordinates so that the lengths are measured from the preceding anchor point. Shifting coordinates to the new anchor, this gives the new initial conditions 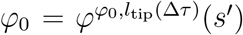 and *l*_0_ = *l*_tip_(Δ*τ*) − *s*′ for the next attachment time. This can be viewed as a discrete-time Markov chain on the states of *l*_tip_ ∈ ℝ+. Generalized into the *n*th step, the process is illustrated in Figure 2 and summarized as follows:

**Figure 2:**
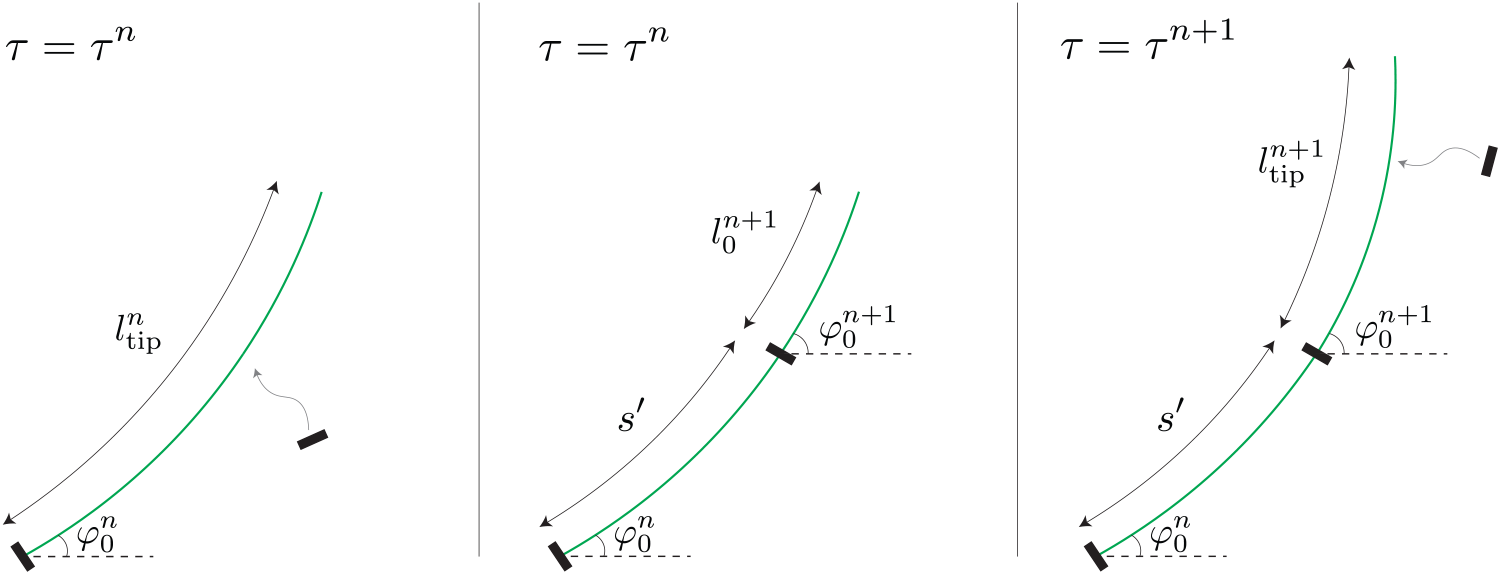
Steps of the anchoring process from the *n*th to *n* + 1th anchor. Left panel: at anchor time *τ*^*n*^, the MT has grown a tip length 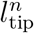. The shape of the MT at this time is determined by the solution to Equation (2) given 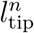 and initial direction of growth 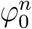 fixed by the preceding anchor. Middle panel: the new anchor position is randomly chosen to be at point *s*′ along the tip, leaving an unanchored length of 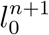. The anchor fixes the angle at position *s*′, and this angle serves as the new initial angle 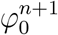 in determining the next tip shape. Right panel: this process is repeated with a new anchoring event at *τ*^*n*+1^ and the corresponding tip length 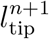.

1. Let *τ*^*n*^ be the time of the *n*^*th*^ anchoring, with tip length 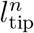 and anchoring angle 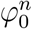 at the *n* − 1^*th*^ anchor. Decide where to anchor uniformly along 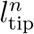.
2. The unanchored segment length is 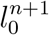 at *τ*^*n*^ with the newest anchoring angle 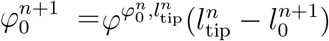
3. Calculate the length that will be grown before next anchoring event. Free tip length is 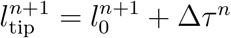

In all, this system is described recursively with compound random variables given by density functions. 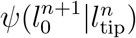 is the probability density of having initial tip length for the *n* + 1^*th*^ anchor, 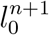, given anchoring at the previous tip length 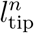. This is uniform. 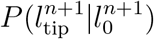 is the density function of having 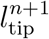 given initial tip length 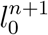 from the previous step. This was found in Equation (6). Together, we have,

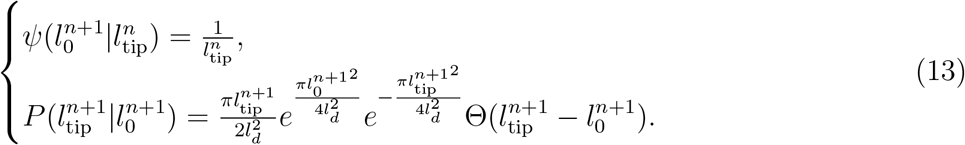

The above describes the chronological steps of anchoring and MT deflection process. Note that the calculation of all anchoring positions and times can be pre-computed. This determines the segments lengths, upon which the deflection *φ* at each segment can be calculated. In this sense, the two processes are decoupled.

### 2.5 Steady State

The stationary average tip length *l*^∗^ may be of interest for estimating regimes which result in buckling. However the relevance of the stationary distribution depends on the convergence rate and length of MT. If the MT is not long enough for the process to converge, this distribution may not be useful. We provide an approximation here, without further analysis.

The two-step process can be reduced to a single step, with the Markov kernel *K*(*l*|*l*^*n*^) describing the probability density of having tip length *l* given *l*^*n*^ previously. The subscript *tip* will be dropped for these variables. To find *K*, we first find the density describing anchoring at point 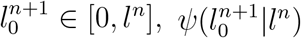. This is uniform. Then we condition the probability of growing to length *l* given initial 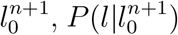, to get 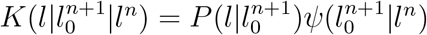. Integrating over the intermediate variable denoted as *l*_0_ rather than 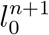,

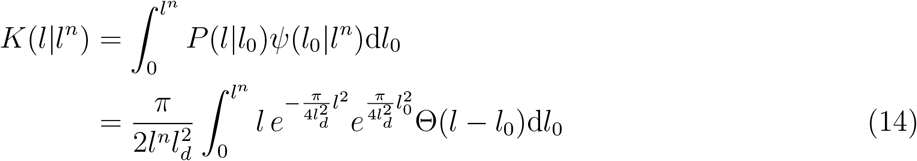

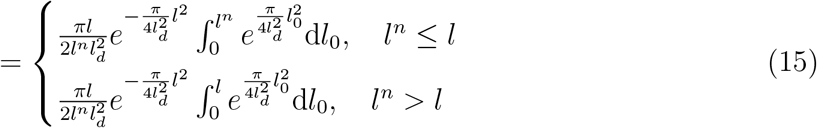

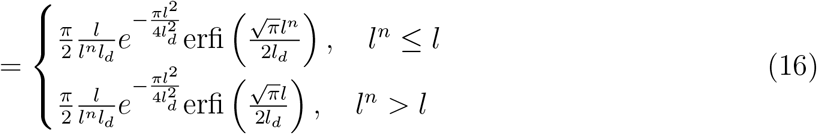

Suppose there exists a stationary distribution *K*_*s*_(*l*). Then, given that this has been reached, the next step will have this same distribution. This is given by the Fredholm integral equation

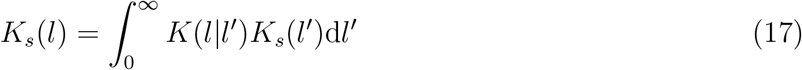

This cannot be solved analytically. Rather, we make a closure approximation that the stationary distribution is approximated by the distribution given by having length *l*^∗^ previously,

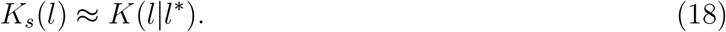

Heuristically, this is true if the stationary distribution is sharply peaked around *l*^∗^. In such case, the previous step, having reached steady state, will sample from *K*_*s*_(*l*) and return values near *l*^∗^. Indeed, in the limit *K*_*s*_(*l*) = *δ*(*l* − *l*^∗^),

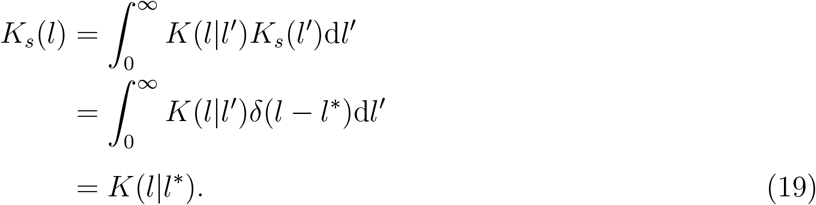

Then one is able to solve for *l*^∗^ with the relation

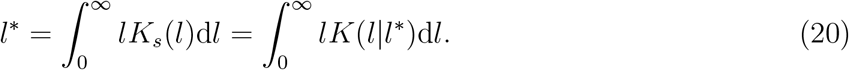

This is confirmed to be accurate numerically in Figure 3 for small values of *l*_*d*_. Larger values of *l*_*d*_ requires more a careful implementation of the numerical root-solving for Equation (20), not attempted here. One notes that as *l*_*d*_ increases, the approximation worsens. This is expected because, as *l*_*d*_ increases, the tip length distribution widens - diverging from the delta function distribution limit.

**Figure 3:**
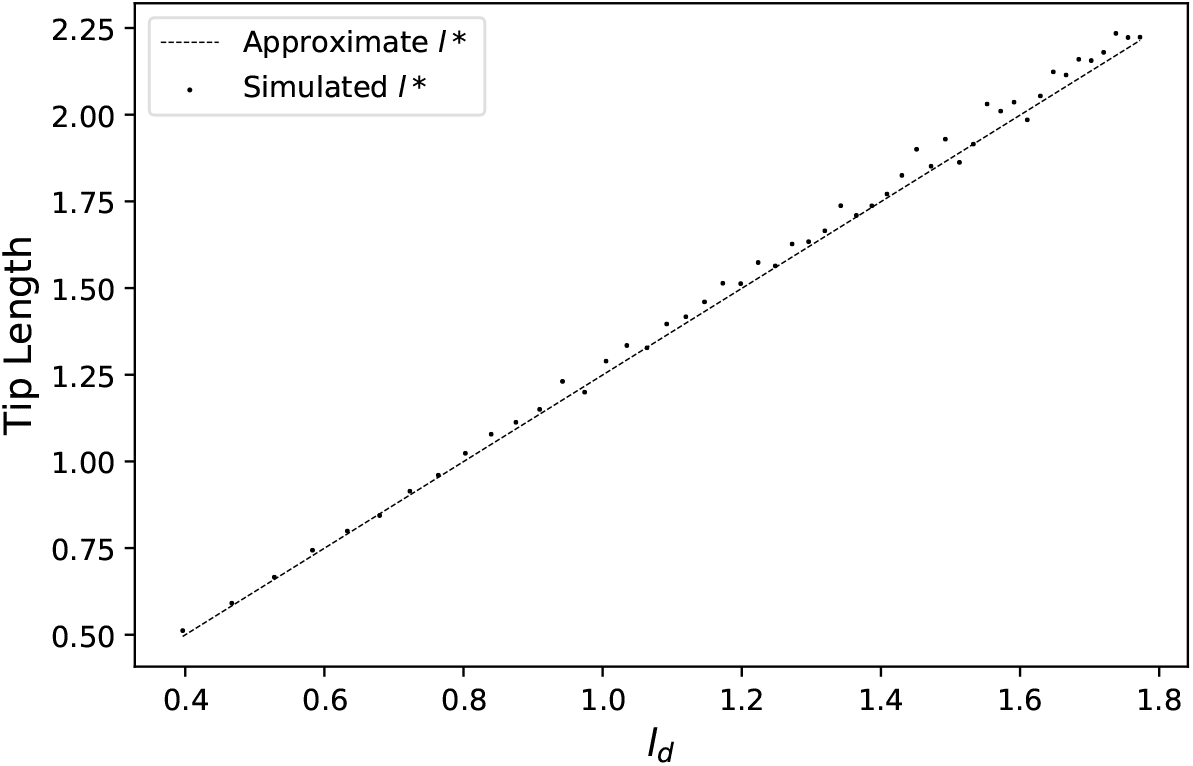
Numerical confirmation of the steady state tip length approximation Equation (20), as a function of small values of *l*_*d*_. Starting with *l*_0_ = 0, 1000 trials were simulated with 40 anchoring steps each. The steady state tip length *l*^∗^ is found by averaging the final tip lengths of the 1000 trials (dotted). The approximate *l*^∗^ was found by numerically solving Equation (20) for *l*^∗^.

### 2.6 Intermediate and Final Positions

As the tip is growing, its position is assumed to be that which instantaneously satisfies the EL equation, 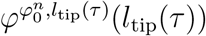. The superscript *n* on *l*_tip_ is dropped to indicate that we are considering times intermediate to anchoring times. This is physically reasonable given the relaxation time of the MTs [13]. The simulation runs until the final generated timestep *τ* ^*N*−1^ *< τ* ^*N*^ = *τ* ^end^ where *τ* ^*N*^ is a user specified value. Thus at *τ* ^*N*^, there is no anchoring event and one must decide what the final position is. This unanchored tip is assumed to be instantaneously in the energy minimizing configuration. Equivalently, it is treated as the end being anchored at *τ* ^*N*^.

Let 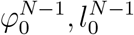 be the last anchored direction and length of free tip after the anchoring, respectively. Since there is no anchoring in (*τ* ^*N*−1^, *τ* ^*N*^], the remaining growth time is Δ*τ* = *τ* ^*N*^ − *τ* ^*N*−1^. Thus, the final angle is 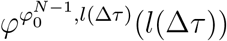.

## 3 Results

We analyze the deflection of MTs in this model in two scenarios. In the first, we calculate the parameter dependence of the distribution of deflection after a single random anchoring event. Here, the MT length is sampled according to the probability distribution in Equation (6) with *l*_0_ = 0. This is a simple case where one can better understand how deflection depends on the tip length and anchoring positions. In the next scenario, we examine the deflection after the MT grows a prescribed length, allowing for any number of anchoring events to occur during growth. This is a more realistic case because MT lengths are generally constrained by the presence of obstacles and/or spontaneous catastrophe.

### 3.1 Analysis of a Single Anchoring Event

We consider the deflection of a MT that nucleates (*l*_0_ = 0) at an initial angle *φ*_0_ and grows until the attachment of the first anchor. Let *φ*^*l*^(*s*) describe the shape for a MT of length *l*, with initial angle satisfying *φ*^*l*^(0) = *φ*_0_.

In the stochastic model, the tip length upon anchoring *l* is sampled from the distribution given in Equation (6). To be concise, we hereafter refer to the tip length upon anchoring as *anchoring length* and emphasize that this is distinct from the *anchor position*. With an anchoring length *l*, the anchor position *s*′ is uniformly distributed over [0, *l*]. Deflections for the stochastic model are found by the difference between *φ*_0_ and *φ*^*l*^(*s*′). Note that *s*′ ≤ *l* and what happens to the shape of the MT between *s*′ and *l* is only determined after a subsequent anchoring. Because we are considering deflection as of a single anchoring event and the deflection calculated as above only accounts for deflection up to the anchor point, it should be thought of as a lower bound for the deflection all the way at *l*.

To explore the parameter dependence of deflection in a simple case, we first consider *φ*_0_ = 0, for which deflection is a result of buckling. The onset of buckling depends only on the anchoring length, and not the subsequent anchor position. We compare the stochastic model, in which the anchoring length is determined by the *l*_*d*_-parameterized distribution given in Equation (6), with a deterministic model, in which the anchoring length is given by the mean of that distribution. To determine the mean anchoring length 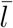 when *l*_0_ = 0, we start by finding the change in length for any *l*_0_ and *φ*_0_ (which is actually independent of *φ*_0_):

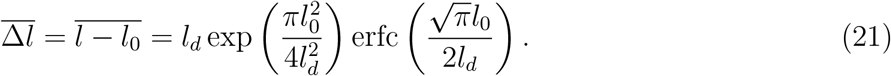

When *l*_0_ = 0, this simplifies to,

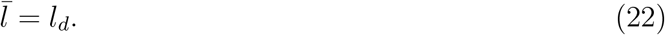

Thus, the non-dimensional length parameter 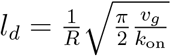 (see Equation (7)) is precisely the mean anchoring length when *l*_0_ = 0. Once an anchoring length *l* is determined, the EL Equation (2) describes the shape the MT takes with the specification of *φ*_0_. For *φ*_0_ = 0, buckling occurs when 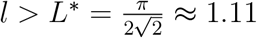 (see Section 2.1). In the deterministic model, *l* is always *l*_*d*_. In the stochastic model, *l* is sampled from the distribution given by Equation (6) and the probability of sampling an anchoring length that leads to buckling can be written as *P* (*l > L*^∗^| *l*_0_ = 0), which implicitly depends of *l*_*d*_ via Equation (6).

Unlike in the deterministic model, when anchoring length is stochastic, buckling can occur even when *l*_*d*_ *< L*^∗^ and can fail to occur even when *l*_*d*_ *> L*^∗^. This is illustrated in Figure 4 where the buckling probability in the deterministic model (blue curve) is either 0 or 1 and the stochastic buckling probability is shown as the black curve. We superimpose the distribution of expected experimental *l*_*d*_ values, to see the overlap between expected mean tip lengths and buckling in both deterministic and stochastic cases. This distribution is generated using values of *v*_*g*_ = 3.5 min^−1^*μ*m and *k*_on_ = 0.34 min^−1^*μ*m^−1^ taken from [11] and *R* (used for non-dimensionalization) distributed normally according to measured values from [4]. Distributions with *k*_on_ an order of magnitude higher and lower are also shown to illustrate sensitivity. Taking the middle distribution as the best estimate of reality, we can compare bucking in deterministic and stochastic models. Because there are more MTs in the middle of the distribution with anchoring lengths just below *L*^∗^ than just above, the net effect is that the stochastic model ought to demonstrate more buckling. For the higher *k*_on_ value, the opposite is true – nearly all estimates of *l*_*d*_ are in the interval between *L*^∗^ and the point at which blue and black curves are indistinguishable. For the lower *k*_on_ value, both models predict essentially no buckling so essentially no difference (but slightly more stochastic buckling if anything).

**Figure 4:**
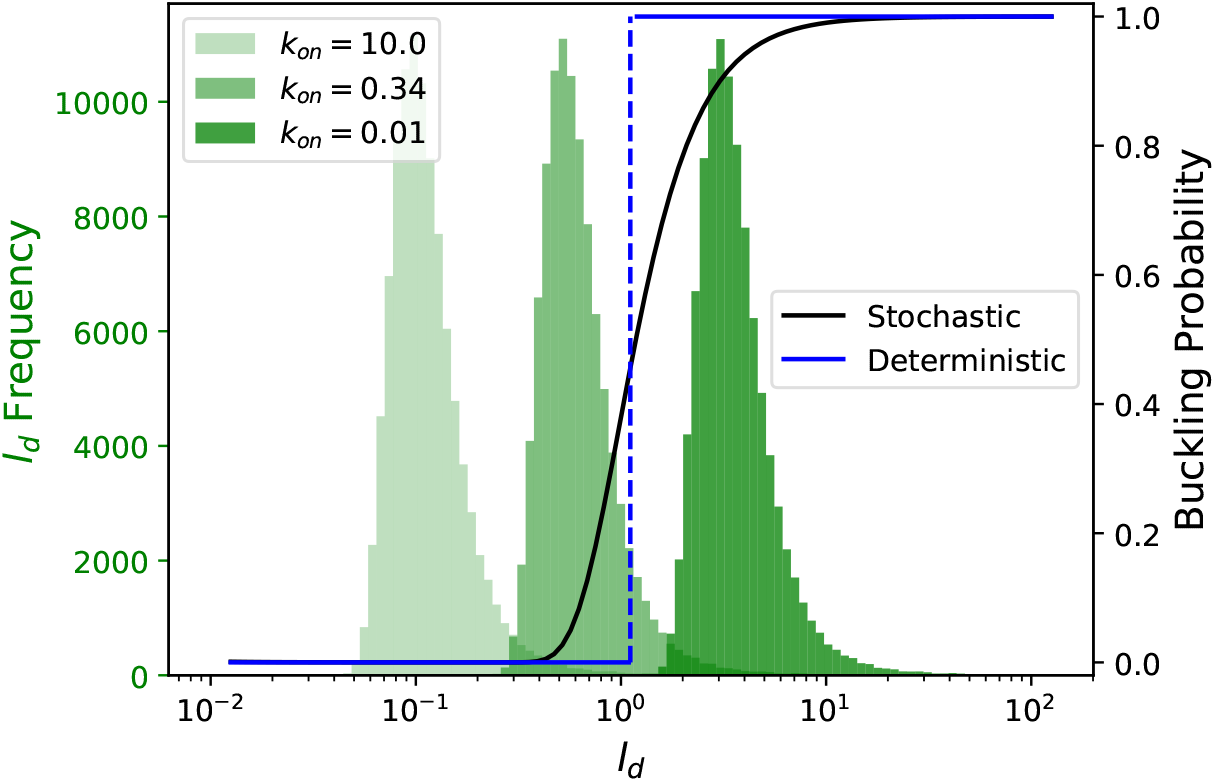
Buckling probability as a function of average tip length *l*_*d*_ with initial length *l*_0_ = 0, as determined from Equation (6). In the deterministic case, the anchoring length is approximated by the mean of the anchoring length distribution from the stochastic model, which in the case of *l*_0_ = 0 considered here is 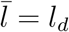. Superimposed is a histogram of *l*_*d*_ estimates generated from *v*_*g*_ and *R* data found in literature, with multiple values of *k*_on_ in units min^−1^*μ*^−1^ (the middle one being from experiment). Depending on *k*_on_, there are varying portions of the population where buckling is significant in both deterministic and stochastic cases.

While buckling indicates whether or not deflection occurs, we also compare the extent of deflection for the deterministic and stochastic cases with a small initial angle *φ*_0_ = 0.01. To do this, it is necessary to consider the resultant shape of the MT. Thus, we must specify further the anchoring position of the above deterministic model. In the deterministic case, we choose to have anchoring happen at the end of the MT which is at *s* = *l*_*d*_. Thus, the end-point deflection is determined by the difference between *φ*_0_ and 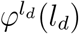. Figure 5 shows deflection values upon anchoring, given *φ*_0_ = 0.01. Superimposed is a histogram of expected *l*_*d*_ values generated with the same data set as in Figure 4. In general, the stochastic deflection has a large variance due to the uniform sampling of the anchor position. One sees that, for *l*_*d*_ small, the stochastic anchoring may be significantly larger than the end-point case. This is because, for small *l*_*d*_, the deterministic anchoring results in a fixed curve of short length whereas the stochastic model may sample larger lengths. MT paths of larger lengths have significant deflection even for short anchor positions *s*′ along its length. Here, 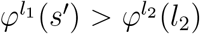 for *l*_1_ *> l*_2_ and *l*_2_ small, even if *s*′ *< l*_2_. Therefore, the sampling of short *s*′ in the stochastic model may still have more deflection. However, as *l*_*d*_ increases, both models have large MT lengths. With larger MT lengths, the curves converge so that 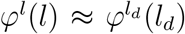 even if *l > l*_*d*_ where both are large. Therefore, the stochastic sampling of short arc positions serves to lower the average deflection. These considerations illustrate the interplay between MT length and anchoring position when determining the deflection, as can be seen in the bottom of Figure 5.

**Figure 5:**
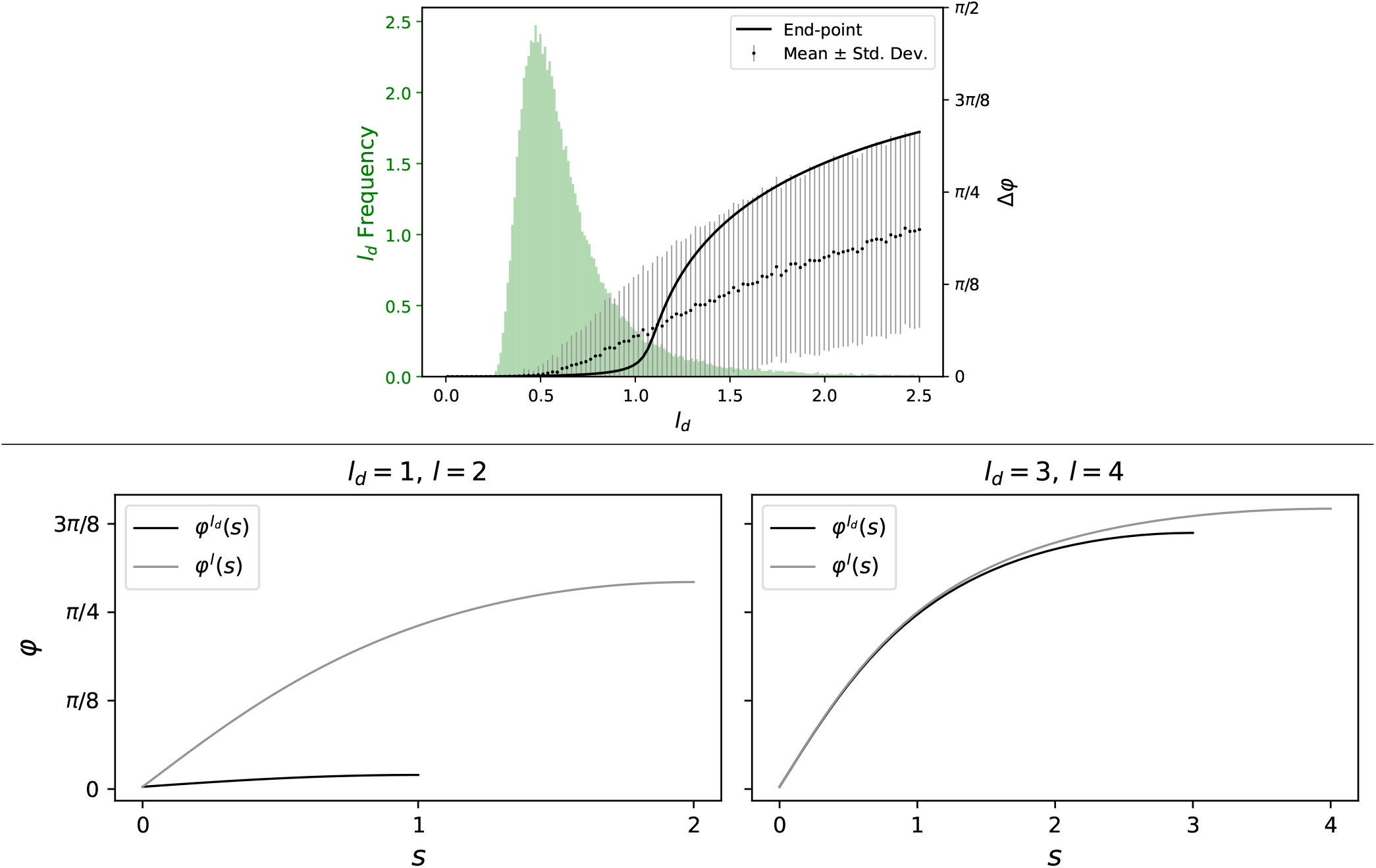
Deflection of a MT as compared to a deterministic model. Top: a comparison of deflection Δ*φ* (mean - dots, standard deviation - bars) of the stochastic model with end-point anchoring given initial tip length *l*_0_ = 0 and angle *φ*_0_ = 0.01. Superimposed is a distribution of average tip lengths 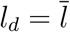 with experimentally found values of *v*_*g*_, *k*_on_, *R*. The stochastic deflection values Δ*φ* are found by difference between *φ*_0_ and *φ*^*l*^(*s*′) where *φ*^*l*^ is the path for MT length *l* sampled from Equation (6) and *s*′ is the location of the first anchor. The deterministic end-point deflection is determined by the difference between *φ*_0_ and 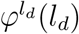 where *l*_*d*_ is the MT length prior to the first anchoring event. Stochastic anchoring generally leads to larger deflection for small *l*_*d*_. Bottom: an illustration of the interplay between MT length and anchoring position in determining deflection. On the left, *l*_*d*_ is small and the deterministic deflection is found from 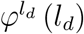. However, the stochastic model may sample lengths *l > l*_*d*_ where 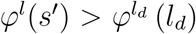 even for *s*′ *< l*_*d*_. The case for larger *l*_*d*_ is shown on the right, where larger sampled MT lengths no longer have significantly larger deflection for any *s*′ value. As a result, sampling smaller *s*′ lowers the average deflection as compared to 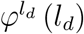.

#### 3.2 l_d_ Controls Deflection

We now analyze the case where the MT grows for a specified length, with possibly many anchoring events. The statistical data of deflection is found through simulations of anchoring and the accompanying EL paths. There are two important aspects: the uninterrupted final length of MT growth *l*_*f*_ and the parameter *l*_*d*_. Additionally, we are interested in the dependence of deflection of the initial angle *φ*_0_. In all, the parameters of interest are *φ*_0_, *l*_*f*_, and *l*_*d*_. Deflection is measured as the difference Δ*φ* = *φ*(*l*_*f*_) − *φ*(0). First, histograms of deflection with fixed *l*_*f*_ and *l*_*d*_ are shown in Figure 6 with different values of *φ*_0_. A density plot composed of histogram data from a range of *φ*_0_ values is shown in the same figure. The maximum elastic deflection Δ*φ*_*max*_ is given by the EL solution with an unsegmented MT of length *l*_*f*_ (black).

**Figure 6:**
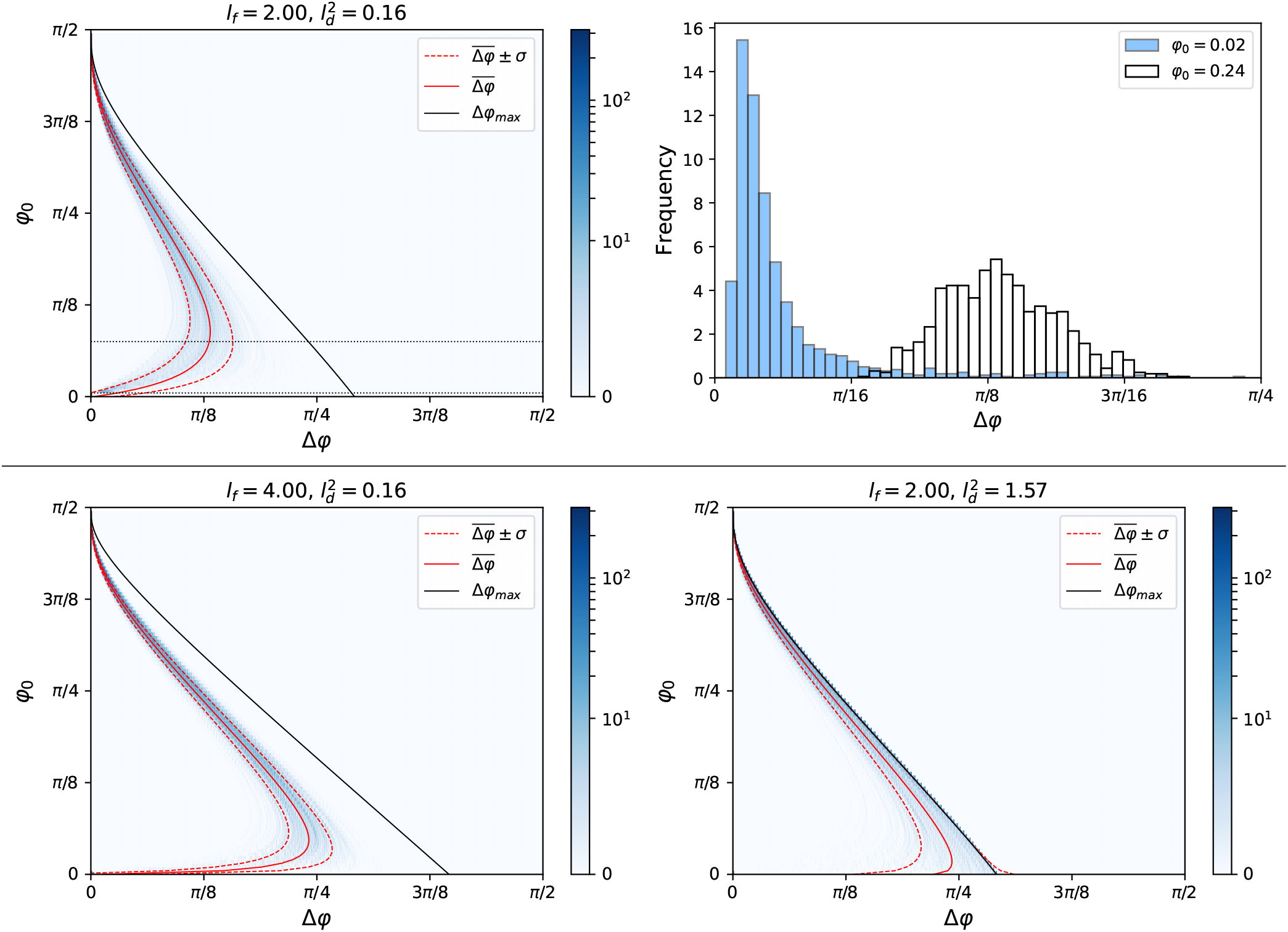
Deflection dependence on the initial angle *φ*_0_. Top right: histograms of total angular displacement Δ*φ* with fixed final MT length *l*_*f*_ = 2 and anchoring length parameter 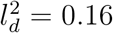. A different *φ*_0_ value is chosen for each. 500 trajectories are sampled for each histogram. Top left: a density plot in which each row represents a normalized histogram for a given *φ*_0_. Dotted black horizontal lines correspond to the histograms for *φ*_0_ = 0.02 and *φ*_0_ = 0.29 shown at the top right. The intensity indicates the densities. Mean deflection (solid red) and values one standard deviation away (dotted red) are also displayed. The 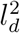 value used is small in the sense that few segments are longer than *l*_*crit*_. This means trajectories are close to geodesics and the mean deflection goes to zero as *φ*_0_ → 0 (segments do not buckle). Bottom: density plots with differing *l*_*f*_ and *l*_*d*_ values. Buckling is seen in the left where 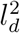 is increased, resulting in large Δ*φ* for *φ*_0_ → 0.

The general behaviour for nearly longitudinal MTs *φ*_0_ → *π/*2 is expected: starting nearly vertical, these MTs do not deflect much as they asymptotically approach *φ* = *π/*2. Of particular interest are the nearly transverse MTs, *φ*_0_ ≈ 0. These MTs exist near the buckling regime and have the potential to deflect greatly. For small *l*_*d*_ (two left panels of Figure 6), each MT segment is small, reducing the deflection toward zero, in keeping with the behaviour of the limiting geodesic case. In particular, in the *φ*_0_ = 0 case, the MTs tend to remain transverse (no buckling). Increasing the total length *l*_*f*_ generally serves to stretch the plots horizontally because an increase in length generally means more segments of a similar length. Consistent with the horizontal stretch, in the case of small *l*_*d*_, the zero deflection of transverse MTs is maintained. However, it also means an increased potential for longer segments, especially in the case of large *l*_*d*_. As seen in the bottom-right plot of Figure 6, increasing 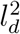 does impact the *φ*_0_ = 0 case by allowing some segments to be longer than *L*^∗^ (buckling). Even for |*φ*_0_| = 0 but small, the deflection can be reduced with small 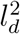. However, this is only valid for a small neighbourhood of *φ* = 0 before deflection becomes significant. This is shown in the upper histograms of Figure 6.

Next, *φ*_0_ and *l*_*f*_ are fixed, and *l*_*d*_ is varied. We examine the transverse trajectories closer in Figure 7. In the top left panel, for *φ*_0_ = 0 there is a clear transition from zero to non-zero deflection. As *l*_*d*_ is increased, segments lengths start to exceed *L*^∗^, initially just outliers and eventually all segments for *l*_*d*_ sufficiently large. Upon the onset of buckling of a segment, subsequent segments further deflect from *φ* = 0. This results in a transition from predominantly zero deflection to large deflection, with a transient bimodal distribution in between. For *φ*_0_ = 0 and larger *l*_*f*_ (not shown), the transition occurs for smaller values of *l*_*d*_. This is because a longer MT allows for more anchoring events and longer segments. In general, as *l*_*f*_ → ∞, we must let *l*_*d*_ → 0 to prevent buckling. The behaviour for different values of *l*_*f*_ and *φ*_0_ ≠ 0 are also shown in Figure 7 (bottom panels). In these cases, there is a unimodal distribution which shifts to the EL limit as *l*_*d*_ is increased. Because geodesic paths given nonzero *φ*_0_ are not stationary solutions of the EL, deflection is unavoidable. Depending on the proximity to *φ*_0_ = 0, deflection may be mitigated.

**Figure 7:**
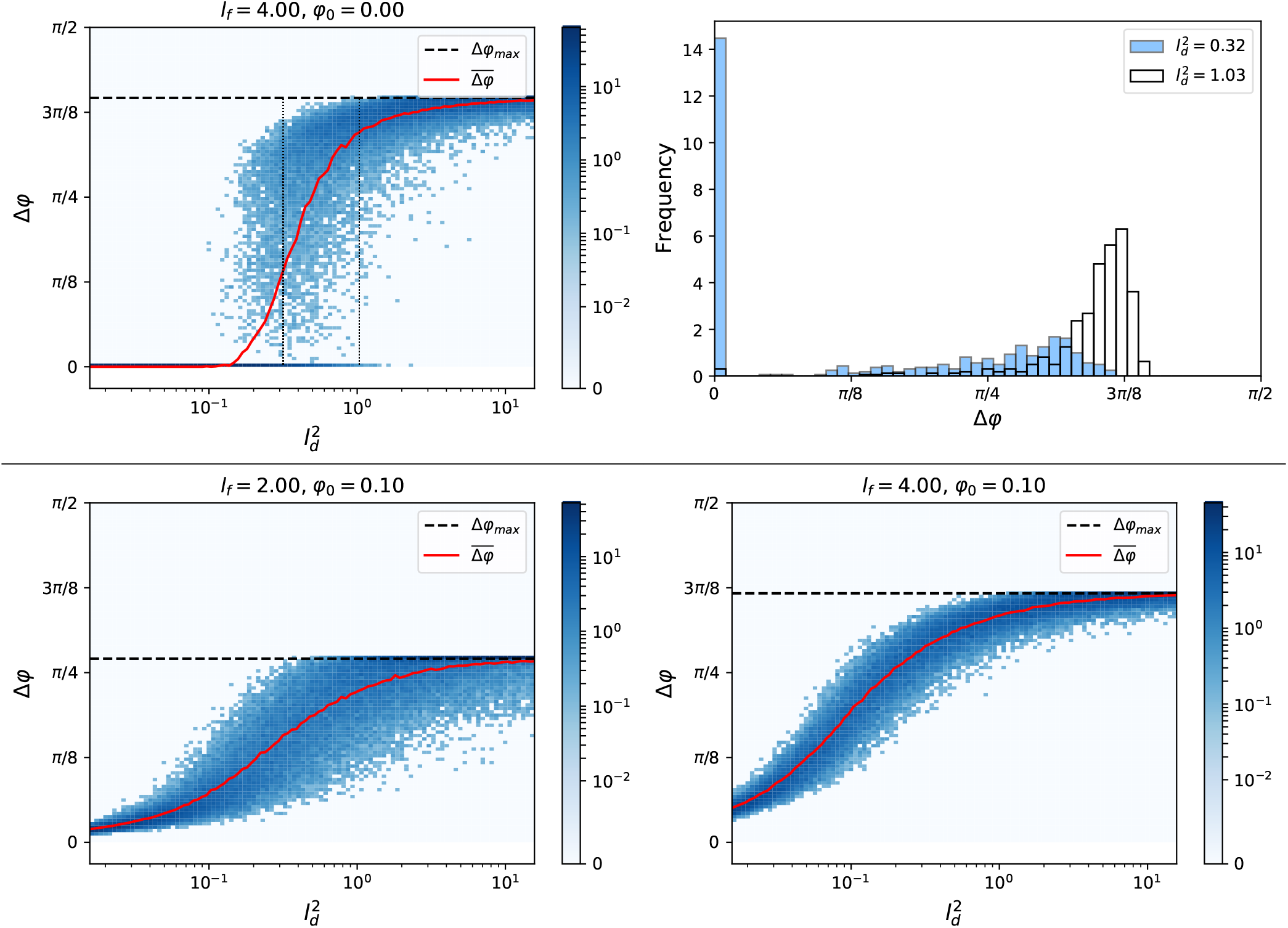
Deflection dependence on the length parameter *l*_*d*_. Top-right: histograms of total angular displacement Δ*φ* with fixed final MT length *l*_*f*_ = 4 and initial angle *φ*_0_ = 0. A different anchoring length parameter *l*_*d*_ is chosen for each. 500 trajectories are sampled for each histogram. Top-left: density plot in which each column represents a normalized histogram for a given *l*_*d*_ as on the right. Dotted black vertical lines correspond to the histograms of select 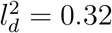 and 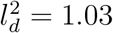 on the right. The intensity indicates the probability densities. Mean deflection (solid red) is also displayed. Bottom: density plots for different values of *l*_*f*_ and *φ*_0_.

## 4 Discussion

Previous modelling works exploring plant cell cortex MT array organization have reproduced the *in vivo* observation of transverse coiling of MTs during the elongation phase, while assuming MTs grow along geodesics. In the case of planes representing faces of polyhedra that approximate cell geometries [7, 4], these are straight lines. In the case of cylinders [5, 6], these are helices. The use of geodesics is traditionally justified as the limit as inter-anchoring distances become small. On the other extreme, the shape of individual trajectories without anchoring between segments have been modelled as thin elastic rods on cylinders, seeking to minimize a curvature energy functional. The results show longitudinal trajectories being favoured, consistent with *in vitro* experiments where MTs were not fully constrained to the surface [9]. A more recent study [10] approximating MT anchoring (also on a cylinder) along uniform segment lengths has found that the trajectories minimizing curvature may deflect significantly from geodesics. In particular, it has been shown that buckling occurs for large enough lengths between anchoring points.

The present work adapts the same elastic rod model, with the addition of non-uniform segment lengths between anchoring. Assuming a uniform rate of anchoring and MT growth, we have proposed a minimal model of the anchoring process and analyzed the resultant distribution of MT segments. With the exception of a few relaxed trajectories corresponding to geodesics, non-zero segment lengths must result in non-zero deflection. Trajectories become biased toward the direction of minimal curvature. On the cylinder, this corresponds to *φ* = ±*π/*2. Unlike the previous work [10], in the present model, deflection may be tuned by modulation of the anchoring process.

We first analyze the deflection in the simple case of a single anchoring event. This case illustrates the interplay between MT length grown prior to anchoring and anchoring position. A stochastic model allows for the possibility of longer tip lengths as compared to when the average length is used. Resulting, the deflection could be large even if the anchoring position is far behind the very tip. This is especially prevalent in the cases of small average tip lengths.

Next, we analyze the model in the more realistic case where the MT grows a prescribed length with possibly many anchoring events. Through the recursive nature of this process, the effects observed in the above case are compounded and difficult to describe analytically or compare to a deterministic analogue. As a result, we more thoroughly examine the dependence of this model on the physical parameters through simulation. Through decreasing the parameter *l*_*d*_, the anchoring distances are decreased and thus minimization of curvature occurs locally. Of particular interest to the question of array organization are the transverse trajectories (*φ*_0_ = 0) which exhibit buckling and those that are nearly transverse (*φ*_0_ ≈ 0) which are generically the ones found in “transverse” arrays and can suffer from comparable deflection to those in the buckling regime. Indeed, it is possible for trajectories in which *φ*_0_ ≈ 0 to maintain small deflections for large MT lengths when anchor density is high. However, assuming nucleation at uniformly random directions, most MTs will not start with *φ*_0_ ≈ 0. For MTs with *φ*_0_ in the range around the peak in Figure 6, deflection can be significant. As a result, most trajectories—if not longitudinal already—are significantly biased toward the longitudinal direction, with a small population maintaining transverse or near-transverse orientations. The question remains (for future work) as to whether these near-transverse MTs are sufficient to produce the observed transverse arrays found in the elongation phase of cells [14]. The mechanically-induced deflection phenomenon complicates and makes this question harder to answer.

In this model and our present analysis of it, array alignment is shown to depend on *l*_*f*_ and *l*_*d*_. This dependence makes the system sensitive to the initial configuration of MTs and nucleation rate. An initially sparse array will allow for longer MTs (larger *l*_*f*_) and thereby more deflection whereas a denser array will suffer less from the mechanically-induced deflections that are the focus of our work. The initial MT arrays are important scaffolding for subsequent MTs to zipper onto and create self-sustaining bundles.

The existence of transverse MTs relies on *l*_*d*_ being sufficiently small. Because 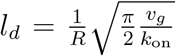, cell regulation could accomplish this through control of the MT growth rate (*v*_*g*_) and/or the rate of anchoring protein attachment (*k*_on_). Macro-scale mechanisms such as edge-induced catastrophe (when MTs reach the edges of cell faces [4]) may not be sufficient in promoting transverse arrays if this initial population of transverse MTs is too small. On the other hand, deflection effects are likely not important once dense arrays are established: *l*_*f*_ and thus deflection becomes very small due to close proximity of MTs. In addition, this may be a mechanism for array-reorientation: starting with transverse arrays, de-polymerization and subsequent lack of anchoring results in large *l*_*d*_, allowing MT trajectories to passively deflect into longitudinal arrays.

The cylindrical geometry provided a simple geometry in which one can investigate the transition between the energetically stable transverse segments and the onset of buckling; one question is whether these results may be applied to other geometries found in cells. This is expected to hold in other geometries which exhibit bifurcations: buckling is delayed by small *l*_*d*_. Stated another way, these trajectories may stay geodesic. On the other hand, for any trajectory in which the minimizer is not geodesic, deflection must occur regardless of the finite anchoring. The determination of minimizers, referred to as relaxed elastics, on other geometries has been studied theoretically by others. In fact, Manning et al. independently derive the equations on a cylinder through a different method, in the context of DNA coiling [15]. Their work can also be applied in this context to surfaces of revolution [16]: *on a surface of revolution, any arc on a meridian is a relaxed elastic line. An arc on a circle of latitude is a relaxed elastic line if and only if the circle of latitude is a geodesic*. Consider an ellipsoid of revolution. It is known (c.f. Clairaut’s relation [17]) that no arc of latitude, other than on the equator, is geodesic. Therefore, purely transverse paths outside the equator are not stationary paths of the curvature functional. Both geodesics and relaxed elastics cannot be transverse in this case. This is in contrast to the cylindrical case where any purely transverse trajectory along the long axis may persist.

Recent experiments on protoplasts approximating ellipsoids show that MTs tend to transiently align with directions of maximum tension upon changes in internal pressure [18]. This would seem to be in contradiction to the longitudinal bias found here. However these experiments analyze trajectories after relatively large time steps that may not capture active MT alignment. It is not conclusive that growing MTs actively align towards this direction. Supposing this does not occur, it is still possible to explain the prevalence of transverse MTs. If tension biases the direction of nucleation and *l*_*d*_ is small, this could result in a net bias to transverse directions. This explanation could be compatible to the proposed model in which tension affects the stability of MTs [12], thereby creating an effective directional nucleation. This could be subject to further investigation.

## Notes

### Competing Interest Statement

The authors have declared no competing interest.

